# Dynamic suspension culture enhances scalable maturation of hiPSC-derived cartilage organoids for regenerative medicine

**DOI:** 10.64898/2026.06.15.732328

**Authors:** Giorgia Mazzini, Evelyn Houtman, Marcella van Hoolwerff, Merel Janssen, Sana Sayedipour, Ghazaleh Hajmousa, Roxanne E. Kieltyka, Rachid R. Mahdad, Yolande F.M. Ramos, Ingrid Meulenbelt

## Abstract

**Background:** Cartilage tissue engineering requires scalable culture strategies to produce high-quality organoids. Human induced pluripotent stem cells (hiPSCs) provide a renewable source of chondrogenic cells. However, conventional static 3D culture limits tissue maturation, reproducibility, and scalability. Dynamic culture systems may help overcome these limitations, although their application for hiPSC-derived cartilage maturation remains poorly explored.

**Methods:** In this study, we established and validated a dynamic suspension bioreactor culture platform (CERO, OLS) for scalable maturation of hiPSC-derived chondroprogenitor cells (hiCPCs) into cartilage organoids populated by biomimetic human induced chondrocytes (hiCHOs). Key culture parameters, including aggregate preparation strategy, agitation speed, and maturation duration, were systematically evaluated. Cartilage maturation under dynamic and conventional static culture conditions was assessed by histology and immunohistochemistry, biochemical assays, organoid size measurements, and gene expression (RT-qPCR). In addition, the functional integration of optimized organoids was evaluated in a human osteochondral explant model.

**Results:** Pre-formed manually picked hiCPC aggregates showed improved cartilage formation compared with single-cell seeding or pelleted aggregates in the bioreactor. Dynamic suspension culture promoted increased construct growth, enhanced ECM deposition, and a more favourable cartilage-associated molecular phenotype compared with static culture. HiCHO organoids matured under dynamic suspension conditions displayed increased sulphated glycosaminoglycan and proteoglycan deposition together with higher expression of cartilage-associated genes *ACAN*, *COMP*, *MGP*, and *COL2A1*. Although prolonged static maturation alone supported continued cartilage development, introducing dynamic suspension culture during later maturation stages further reinforced favourable molecular and matrix-associated features. Importantly, hiCHO organoids generated under optimized dynamic culture conditions successfully filled human cartilage defects and established matrix continuity with surrounding native tissue in a human osteochondral *ex vivo* explant model.

**Conclusions:** This study shows that dynamic suspension culture is an effective and scalable strategy for maturation of hiPSC-derived cartilage organoids. Consequently, this approach supports reproducible neo-cartilage production and allows functional testing in human tissue models. These findings support the use of dynamic culture systems for cartilage repair and *in vitro/ex vivo* cartilage research.

## Background

Osteoarthritis (OA) is a degenerative joint disease caused by cartilage breakdown, leading to pain, stiffness, and reduced mobility (1, 2). Cartilage consists of chondrocytes that produce and maintain dense extracellular matrix (ECM). However, during OA this mechanism fails, leading to cartilage degeneration and remodeling of subchondral bone .

Autologous cartilage implantation (ACI) is a promising regenerative therapy for cartilage repair (3, 4), however, isolating human primary articular chondrocytes (hPACs) from patients is invasive, the availability of cells is restricted in severe osteoarthritic joints, and the chondrogenicity of these cells decline in culture . These challenges may be addressed by human induced pluripotent stem cell (hiPSC) technology. hiPSCs are created by reprogramming somatic cells into stem cells that have broad differentiation potential and proliferative capacity, offering a sustainable, non-invasive source for cartilage generation (5). It has been shown that hiPSCs differentiated into chondroprogenitor cells (hiCPCs) and subsequently into chondrocyte-like organoids (hiCHOs) closely resemble hPACs when cultured in static suspension systems (6–8). However, this approach remains labour-intensive and time-consuming, highlighting the need for more efficient and scalable culture strategies (8, 9).

Bioreactor technologies provide controlled and physiologically relevant environments for tissue generation, enhancing scalable matrix deposition and mechanical resilience (9, 10) . Various platforms, including rotating wall vessels, spinner flasks, and perfusion systems, have been applied to cartilage engineering (11–13), though not yet to hiPSC-derived cartilage. The 3D dynamic suspension culture using a bioreactor represents a promising option, as it has already supported scalable differentiation of hiPSC-derived hepatocytes, microglia, cardiomyocytes, and macrophages (14). Moreover, dynamic suspension culture of murine iPSCs has been shown to accelerate chondrogenesis (11, 12) and increase proliferation, suggesting that suspension cultures in bioreactors could be well suited for hiPSC-derived neo-cartilage maturation.

By integrating stem cell biology with dynamic bioreactor engineering, we sought to establish a scalable and reproducible platform for maturation of hiPSC-derived cartilage organoids suitable for translational cartilage research and regenerative medicine. To achieve this, we employed a dynamic suspension bioreactor to support scalable and high-quality cartilage maturation from hiCPCs. Finally, to assess the potential for translational relevance of these constructs, we evaluated their capacity to integrate into human osteochondral tissue using an established human joint tissue *ex vivo* explant model obtained from OA patients undergoing joint replacement surgeries (15, 16). Establishing such a workflow contributes to advance systems and engraftment strategies towards effective stem cell–based regenerative therapies for osteoarthritis and related cartilage disorders.

## Materials and Methods

### Sample description and ethics approval

Primary chondrocytes and osteochondral explants were obtained from knee joints of osteoarthritis (OA) patients undergoing total joint replacement surgery as part of the Research in Articular Osteoarthritis Cartilage (RAAK) study (17). Ethical approval was granted by the Medical Ethics Committee of the Leiden University Medical Center (P08.239 / P19.013), and informed consent was obtained from the donor. Donor characteristics are provided in **Supplementary Table S8.** A control hiPSC line were used in the current study LUMC0004iCTRL10 (“LUMC0004” here). LUMC0004 cells were generated from skin fibroblasts of a male by the LUMC iPSC core facility and registered at the Human Pluripotent Stem Cell Registry. Approval for the generation of hiPSCs from skin fibroblasts of healthy donors is available under number P13.080. LUMC0004 cells were characterized according to pluripotent potential and spontaneous differentiation capacity by the iPSC core facility (18).

### hiPSCs culture and maintenance

hiPSCs were maintained in mTesR Plus medium (STEMCELL Technologies) on VitronectinXF-coated plates (STEMCELL Technologies), at 37 °C in 5 % CO2. Medium was refreshed every two days and at approximately 80 % confluency the cells were passaged using Gentle Cell Dissociation Reagent (STEMCELL Technologies). Mycoplasma testing was performed on a regular basis.

### Stepwise hiPSC differentiation into human induced chondroprogenitor cells (hiCPCs)

Generation of induced chondroprogenitor cells (hiCPCs) was based on a protocol previously described (6, 7, 19). When hiPSCs reached 60 % confluence, the culture medium was switched to mesodermal differentiation (MD) medium, composed of IMDM GlutaMAX (IMDM; Thermo Fisher Scientific) and Ham’s F12 Nutrient Mix (F12; Sigma-Aldrich) with 1 % chemically defined lipid concentrate (Gibco), 1 % insulin/human transferrin/selenous (ITS+; Corning), 0.5 % penicillin-streptomycin (P/S; Gibco), and 450 μM 1-thioglycerol (Sigma-Aldrich). Before induction of anterior primitive streak (day 0), hiPSCs were washed with wash medium (IMDM/F12 and 0.5 % P/S) and then fed with MD medium supplemented with activin A (30 ng/ml; Stemgent), 4 μM CHIR99021 (CHIR; Stemgent), and human fibroblast growth factor (20 ng/ml; FGF-2; R&D Systems) for 24 hours. Subsequently, the cells were washed again with wash medium, and paraxial mesoderm was induced on day 1, by MD medium supplemented with 2 μM SB-505124 (Tocris), 3 μM CHIR, FGF-2 (20 ng/ml), and 4 μM dorsomorphin (Tocris) for 24 hours. Before induction of early somite (day 2), cells were washed with wash medium, and then cells were fed with MD medium supplemented with 2 μM SB-505124, 4 μM dorsomorphin, 1 μM C59 (Cellagen Technology), and 500 nM PD173074 (Tocris) for 24 hours. Subsequently, cells were washed with wash medium, and for induction of sclerotome, cells (days 3 to 5) were fed daily with MD medium supplemented with 2 μM Purmorphamine (Stemgent) and 1 μM C59. To induce chondroprogenitor cells (days 6 to 14), cells were washed briefly with wash medium and fed daily with MD medium supplemented with human bone morphogenetic protein 4 (BMP-4; 20 ng/ml; Miltenyi Biotec).

### Chondrogenic differentiation and maturation in 3D suspension cultures

Chondrogenic differentiation and maturation were performed in three-dimensional (3D) suspension cultures under either static conditions or dynamic rotary suspension using the CERO bioreactor (OMNI Life Science, Germany), as detailed below. For 3D maturation, all samples were cultured in chondrogenic differentiation medium (CDM) consisting of DMEM/F12 + GlutaMAX (Gibco, Thermo Fisher Scientific), insulin–transferrin–selenium (ITS+), MEM non-essential amino acids (NEAA), dexamethasone, penicillin/streptomycin, 2-mercaptoethanol, L-ascorbate, L-proline, and transforming growth factor beta 1 (TGF-β1; Peprotech). Medium supplements were freshly added prior to each medium change, which was performed every two to three days. For dynamic suspension culture in the bioreactor, hiCPC aggregates were washed with phosphate-buffered saline (PBS) and, where indicated, dissociated using TrypLE prior to seeding. Aggregates or cells were transferred into bioreactor tubes containing 10 mL of CDM and cultured under defined rotary suspension conditions. Manufacturer-recommended user program settings applied during the inoculation and culture phases are summarized in **Supplementary Table S1**, unless stated otherwise.

### Optimization of hiCPC inoculation strategy for aggregate formation

Preliminary experiments were performed to identify the most effective inoculation strategy for generating uniform hiCPC aggregates suitable for dynamic suspension culture. Day-14 hiCPCs were initially seeded as single-cell suspensions in the bioreactor at different starting densities, including 1.0 × 10⁶ and 2.0 × 10⁶ cells per tube, to evaluate spontaneous aggregate formation under dynamic suspension conditions. In parallel, pre-formed hiCPC aggregates were evaluated as an alternative inoculation strategy. Aggregate formation efficiency, morphological uniformity, and compatibility with subsequent chondrogenic maturation under dynamic suspension culture were assessed to determine the optimal inoculation approach.

To evaluate whether pelleting could provide a less labor-intensive alternative to manual picking, at day 14 of differentiation, hiCPCs were prepared as either manually picked aggregates or pelleted aggregates. For the pelleting workflow, dissociated hiCPCs (2.5 × 10⁵ cells) were centrifuged at 1640 rpm to form pellets, which were pre-incubated under static conditions for 24 hours to allow re-aggregation. Both picked aggregates and pelleted aggregates were subsequently cultured either in static 15 mL conical tubes or in bioreactor tubes under dynamic suspension conditions. This experiment was designed to determine whether pelleted aggregates could replace manually picked aggregates without compromising aggregate uniformity, size, or chondrogenic matrix formation.

### Optimization of dynamic suspension culture parameters for hiCHOs maturation

Dynamic bioreactor suspension parameters were optimized to promote aggregate maturation and extracellular matrix (ECM) deposition. Variables including rotation speed and culture duration were systematically assessed. Picked aggregates were cultured at 60 or 80 rpm for 2 or 4 weeks in 50 mL conical bioreactor tubes containing chondrogenic medium and compared to the conventional static culture. The optimal conditions identified, 80 rpm for 4 weeks, were subsequently used for all downstream experiments. This optimization step established the baseline settings for reproducible dynamic suspension maturation of hiCPC-derived neo-cartilage organoids.

### Exploring extended maturation using combination of static and dynamic suspension culture

To investigate the effect of prolonged maturation, an independent experiment was performed in which hiCPC-derived aggregates were differentiated for 14 days and subsequently matured under static conditions for 4 weeks. At this stage, hiCHO organoids were either maintained under static culture or transferred to the bioreactor for an additional 2 weeks, resulting in a total culture duration of 6 weeks. Throughout the experiment, all hiCHO constructs were maintained under identical chondrogenic culture conditions. This approach was designed to evaluate whether introducing dynamic suspension culture after initial tissue formation could further influence cartilage maturation, matrix deposition, and tissue organization during extended culture.

### Osteochondral explant cultures

Osteochondral explants were obtained from macroscopically intact regions of the femoral condyle within 2 h after surgery from a donor undergoing total knee replacement and prepared according to a previously described protocol (15, 20). To evaluate different defect-creation strategies for graft engraftment, cartilage defects were generated using either biopsy punches or a mechanical rotary drill. Both approaches produced cylindrical defects of approximately 1 mm in diameter and 1 mm in depth. For engraftment, defects were filled with hiPSC-derived chondrocyte-like organoids (hiCHOs) matured under dynamic suspension culture. Prior to implantation, organoids were histologically evaluated to confirm ECM deposition and stable chondrogenic phenotype. hiCHO organoids were gently press-fit into the defects to ensure stable positioning without damaging the surrounding cartilage. To further stabilize the grafts and reduce micromotion at the graft–host interface, a thin layer of biocompatible hydrogel was applied immediately after implantation, following previously described principles (20). This hydrogel gradually dissolved over several days at 37 °C and served as a temporary fixation layer without remaining in the tissue long-term. Explants were cultured in serum-free chondrogenic differentiation medium for 4 weeks at 37 °C and 5% CO₂, with medium refreshed three times per week. At endpoint, samples were processed for histological assessment of matrix deposition and graft–host integration.

### Neocartilage organoid size estimation

To evaluate neo-cartilage growth, hiCHO organoids were imaged at the time of harvest using a brightfield microscope (Olympus CKX53). A region of interest (ROI) was manually defined around the spherical core of each construct using CellSens software (Olympus), excluding irregular peripheral outgrowths, off-target tissue, and non-cartilaginous borders. The area of the ROI was then measured and used as an estimate of pellet size.

### Sulphated glycosaminoglycan (sGAG) quantification

Matrix production was evaluated by quantifying sGAG content using the 1,9-dimethylmethylene blue (DMMB) assay as previously described by Farndale et al. (21). Two pellets per condition were pooled and digested overnight at 60°C in 200 μl of papain solution (Sigma-Aldrich) and diluted 50-fold in ethylenediaminetetraacetic acid (EDTA) solution prior to the DMMB reaction. Shark chondroitin sulphate (Sigma-Aldrich) served as the reference standard. Absorbance was measured at 525 and 595 nm using a microplate reader (Spectramax iD3, Molecular Devices). To normalize for cell number, sGAG concentrations were corrected for DNA content, measured using the Qubit® 2.0 Fluorometer and the dsDNA High Sensitivity Assay Kit (Invitrogen™). Final values were reported as µg sGAG per µg DNA.

### RNA isolation and gene expression analysis (RT-qPCR)

For RNA isolation, from each condition two pellets were pooled and lysed in 200 µl Trizol reagent (Thermo Fisher Scientific) and homogenized using micro pestles. RNA was extracted with chloroform and purified from the supernatant using the RNeasy Mini kit (Qiagen). RNA concentration was measured using a Nanodrop-1000 photospectrometer (Thermo Scientific). Synthesis of cDNA was performed with 150 ng of total RNA using a First Strand cDNA Synthesis kit according to manufacturer’s protocol (Roche Applied Science). The cDNA was diluted five times and expression levels of seven genes of interest were measured with QuantStudio 6 Real-Time PCR system using FastStart SYBR Green Master reaction mix (Roche Applied Science). Primer sequences are shown in **Supplementary Table S2**. Relative gene expression levels (-ΔCt) were calculated using the average of Ct values of Glyceraldehyde-3-Phosphate Dehydrogenase (*GAPDH*) and and Succinate dehydrogenase A (*SDHA*) as internal control (IC). ΔCt values of the gene of interest (GOI) were calculated as Ct_GOI_ – Ct_ic_. To compare lineage-specific gene expression, the relative contribution of chondrogenic (*COL2A1*), fibrotic (*COL1A1*), and hypertrophic (*COL10A1*) transcripts was estimated by calculating normalized ratios of each gene over the total collagen signal, using log₂-transformed Ct values.

### Histological and immunohistochemical (IHC) analysis

Pellets were fixed overnight in 4 % formaldehyde and stored in 70% ethanol at 4°C. Samples were processed using a Tissue-Tek VIP 5 Jr. processor (Sakura), embedded in paraffin, and sectioned at 5 µm using a HistoCore Multicut microtome (Leica). Slides were dried, deparaffinized, and rehydrated prior to staining. Osteochondral explants were fixed in 4% formaldehyde and decalcified using Mol-Decalcifier (Milestone) for one week at 37°C, dehydrated with an automated tissue processor, and paraffin-embedded. Sections of 5 µm thickness were obtained for all downstream analyses. Cartilage matrix composition was evaluated using 1% Alcian Blue 8-GX (Sigma-Aldrich) to visualize sulphated glycosaminoglycans (sGAGs), counterstained with Nuclear Fast Red (Sigma-Aldrich). Proteoglycan content was assessed with 1% Safranin-O (Sigma-Aldrich), with Fast Green (Sigma-Aldrich) to distinguish non-cartilaginous matrix. For immunohistochemistry, sections were incubated with primary antibodies against Collagen Type II (ab34712, Abcam, 1:250), Collagen Type VI (ab6588, Abcam, 1:250) and Ki-67 (ab92742, Abcam, 1:500) (**Supplementary Table S3**). Staining detection was performed using 3,3′-diaminobenzidine (DAB; Sigma-Aldrich) as the chromogen and hematoxylin (Klinipath or Sigma-Aldrich) as a counterstain to visualize nuclei. Quantification of staining intensity was performed using Fiji/ImageJ (version 1.54p). High resolution images of stained neo-cartilage sections were acquired using a ZEISS Axioscan 7 digital slide scanner and converted to 8-bit grayscale negatives. An identical threshold was applied across all samples to ensure consistency. Tissue regions were manually defined as regions of interest (ROIs) to exclude background, and mean grey values were measured as an indicator of positive staining intensity. Because inverted grayscale images were used, higher grey values corresponded to stronger staining intensity.

### Statistical analysis

Statistical analyses were performed in R version 4.4.1 (R Foundation for Statistical Computing, Vienna, Austria). Data are presented as boxplots showing the median and interquartile range (IQR), with whiskers extending to the most extreme data points within 1.5 × IQR. Individual biological replicates are shown as dots. Differences between experimental groups were assessed using generalized estimating equations (GEEs) (22) implemented in the *geepack* package. Differences were considered statistically significant at P < 0.05. Significance levels are indicated as follows: *P < 0.05, **P < 0.01, and ***P < 0.001.

## Results

### Experimental design overview

A schematic overview of the study design is provided in **Supplementary Figure S1**. The study was designed to establish and validate a scalable dynamic suspension culture platform for the maturation of hiPSC-derived cartilage organoids, benchmark its performance against conventional static culture, and assess its translational potential in a human osteochondral explant model. First, we optimized key bioreactor culture parameters required for robust hiCHO maturation. We then compared the optimized dynamic suspension culture with conventional static culture by assessing organoid growth, ECM deposition, histological features, and chondrogenic gene expression profile. Finally, dynamically matured hiCHO organoids were evaluated in a human osteochondral explant model to assess defect filling and matrix continuity with the surrounding native cartilage.

### Pre-formed hiCPC aggregates are required for robust cartilage organoid maturation under dynamic suspension culture

To establish suitable culture conditions for dynamic hiCHOs maturation, day-14 hiCPCs were first introduced into the bioreactor as single-cell suspensions for 2 weeks. Under these conditions, cells failed to consistently form 3D structure, even after increasing the initial seeding density (data not shown). Likewise, although higher cell numbers promoted partial aggregation and clustering, the resulting hiCHO organoids remained poorly organized and showed minimal ECM deposition. In contrast, when we evaluated the use of manually selected pre-formed hiCPC aggregates, we consistently managed to generate compact hiCHO organoids in the bioreactor (**Supplementary Figure S2**). Alcian Blue staining demonstrated robust sulphated glycosaminoglycan deposition and improved tissue organization compared with hiCHO constructs generated directly from single-cell suspensions. After 2 weeks of maturation, these constructs already appeared chondrogenic, although further maturation appeared possible. To then explore the scalability of the workflow and avoid manual picking of individual hiCPC aggregates, maturation in the bioreactor was compared between manually picked pre-formed hiCPC aggregates and aggregates generated by dissociation and pelleting of day-14 hiCPCs. While pelleted aggregates gave rise to larger organoids (**Supplementary Figure S3B-C, Supplementary Table S4**), histological analyses revealed considerable variability in tissue organization and ECM deposition between hiCHO constructs. In contrast, manually picked aggregates consistently generated compact hiCHO organoids consistently positive for Alcian Blue staining (**Supplementary Figure S3D**). Together, these findings demonstrate that preservation of early hiCPC aggregates architecture is essential for reproducible hiCHO cartilage organoids maturation under dynamic suspension culture. Therefore, manually picked pre-formed hiCPC aggregates were used in all subsequent experiments.

### Optimization of dynamic suspension culture parameters

We next examined whether rotation speed and culture duration could influence hiCHOs maturation in the dynamic bioreactor system. Manually picked pre-formed day-14 hiCPC aggregates were cultured at either 60 or 80 rpm for 2 or 4 weeks. In both conditions, pellet area increased over time, with the largest organoids observed after 4 weeks at 80 rpm (**Supplementary Figure S4A**; **Supplementary Table S5A**). Although both conditions supported cartilage matrix formation, hiCHO organoids matured at 80 rpm displayed stronger and more homogeneous ECM staining compared with those cultured at 60 rpm, as evidenced by positive Alcian Blue staining after 4 weeks of maturation (**Supplementary Figure S4B**). In addition, hiCHO organoids cultured at 80 rpm displayed a reduced presence of putative off-target/immature cells at the outer rim compared to those cultured at lower speed. Quantification confirmed significantly higher matrix-associated staining intensities at 80 rpm (**Supplementary Figure S3C; Supplementary Table S5B**). Collectively, these results demonstrate the ability of the bioreactor system to generate large hiCHO constructs with robust ECM deposition, highlighting the capacity of dynamic suspension culture to support scalable tissue formation. Based on these findings, 80 rpm and 4 weeks of maturation were selected as the dynamic suspension culture condition for the subsequent experiments.

### Comparison of dynamic suspension and static culture conditions for hiCHOs maturation

Next, we compared hiCHOs-made cartilage organoid maturation between conventional static and dynamic suspension culture applying the optimized bioreactor conditions. After 4 weeks, hiCHO-made cartilage organoids formed in dynamic suspension were significantly larger, reaching approximately 4 mm^2^ in size (**Figure 1A**; **Supplementary Table S6A**) and had significant increased sGAG/DNA levels (**Figure 1B**; **Supplementary Table S6B**) as compared to conventional static cultures. Histological analysis with Alcian Blue (**Figure 1C**) and Safranin-O (**Figure 1D**) staining confirmed this increased deposition of sGAG and proteoglycans. Notably, the peripheral region of non-cartilage like tissue observed in hiCHO organoids matured in static cultures was largely absent in those cultured in the bioreactor, suggesting reduced tissue heterogeneity and more consistent cartilage formation throughout the construct.

**Figure 1.**
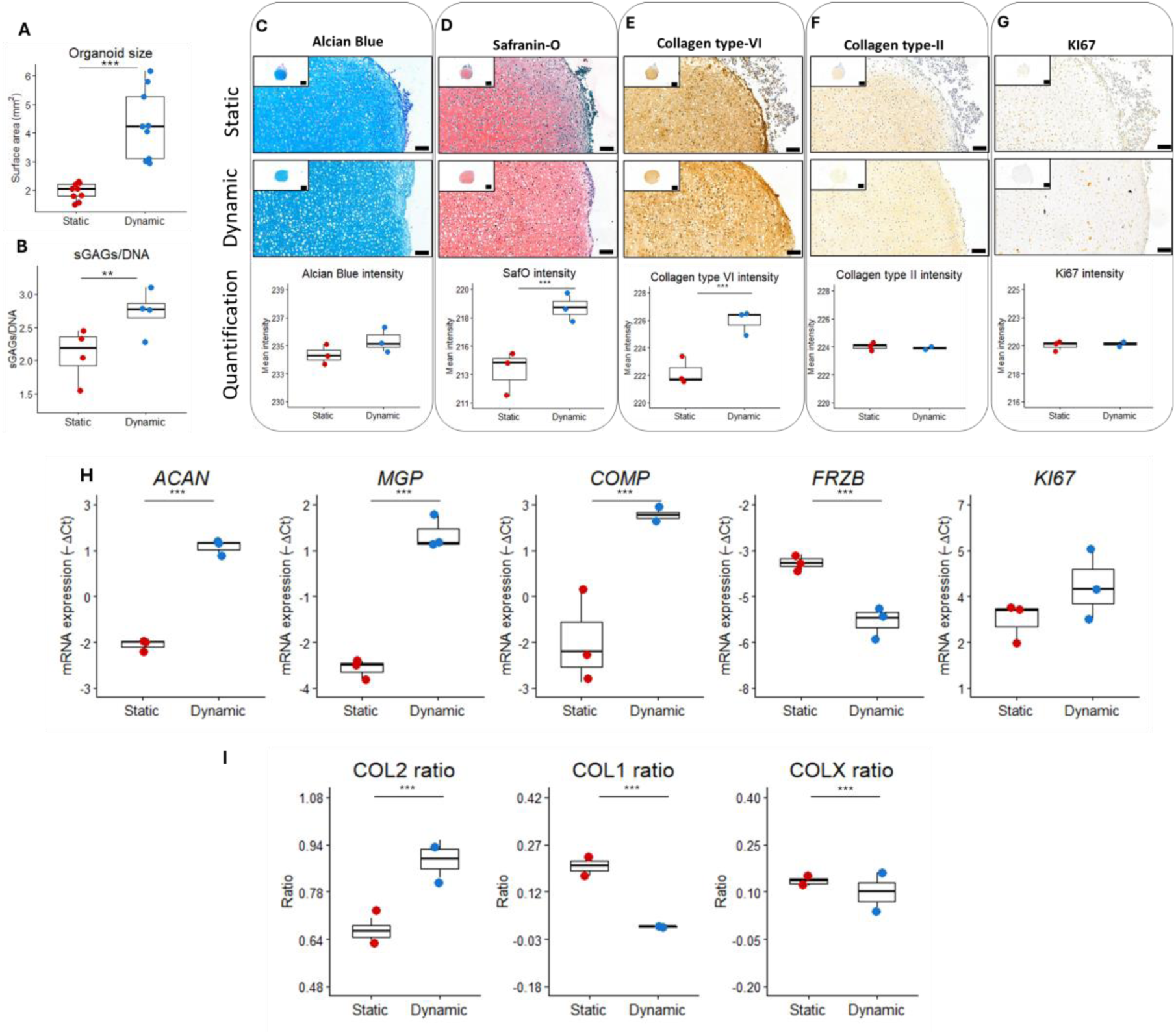
Dynamic suspension culture enhances maturation of hiCHO constructs. (**A**) Scatter boxplot showing hiCHO organoids size between static and dynamic (**B**) Sulphated glycosaminoglycan content normalized to DNA (sGAG/DNA) of hiCHO organoids matured for 4 weeks under static or bioreactor conditions. (**C–G**) Representative histological and immunohistochemical analyses of ECM deposition, cartilage matrix organization, and proliferative activity using Alcian Blue, Safranin-O, collagen type VI, collagen type II, and KI67 staining in W4 hiCHO organoids. Quantification of staining intensity is shown below each representative image. Insets show low and high-magnification overviews (4x, 20x) of the entire constructs. Corresponding images from all biological replicates are provided in **Supplementary Figure S5**. (**H**) Relative mRNA expression levels (−ΔCt) of established chondrogenic markers showing the phenotype of hiCHOs after 4 weeks maturation in both conditions. (**I**) Relative abundance of collagen type II, type I, and type X collagen. Each dot represents an individual construct; bars indicate mean ± SD. Statistical significance was determined using generalized estimating equations (GEE). *P ≤ 0.05, **P ≤ 0.01, ***P ≤ 0.005.

Subsequent immunohistochemistry of collagen type-II (**Figure 1F, Supplementary Table S6C**), displayed an even distribution throughout the ECM in both conditions. Notably was that collagen type-VI deposition appeared significantly increased (**Figure 1E, Supplementary Table S6C**) for hiCHO-formed cartilage organoids in the bioreactor. Moreover, collagen type VI staining was prominently localized around lacunae-like structures, consistent with the formation of a cartilage-specific pericellular matrix (PCM) surrounding individual chondrocytes, referred to as chondrons. Ki-67, a marker for cell proliferation, was not significant different between the conditions (**Figure 1G, Supplementary Table S6C**). To further assess cartilage maturation, we compare cellular phenotypic states of hiCHOs and performed gene expression analysis of some important articular chondrocyte markers. As shown in **Figure 1H** and **Supplementary Table S6D**, hiCHOs matured dynamic suspension showed significant higher expression of *ACAN*, *MGP*, and *COMP.* Additionally (**Figure 1I**; **Supplementary Table S6E**), the abundance of *COL2A1* relative to *COL1A1* and *COL10A1* increased significantly, indicating the formation of hyalin instead of fibrotic or hypertrophic hiPSC-derived cartilage.

### Assessment of prolonged hiCHOs maturation under static and dynamic culture conditions

Having observed the relevance of preformed matrix (**Supplementary Figure S2-3**), we next determined if extending culture for additional 2 weeks could further mature hiCHO-neocartilage with respect to matrix deposition and tissue organization. To this end, after 4 weeks of static hiCHO-maturation a combined static – dynamic culture was compared to hiCHO-organoids maintained in static culture for an additional 2 weeks. As shown in **Figure 2A** (**Supplementary Table S7A**), after 6 weeks of maturation hiCHO made organoids in both conditions were still growing, now all reaching average sizes of approximately 5 mm² while abundantly depositing sGAGs and proteoglycans rich matrix as marked by intense Alcian Blue and Safranin-O staining (**Figure 2B-C**; **Supplementary Table S7B**). Moreover, hiCHO-made organoids abundantly expressed collagen type-II throughout the tissue (**Figure 2E**) whereas collagen type-VI expression was particularly visible around rounded cells and well-defined lacunae, demonstrating that prominent chondron-like structures were formed. The latter particularly in the static cultures (**Figure 2D**). While small regions of non-cartilage tissue were still observed in some hiCHO organoids after prolonged culture (**Supplementary Figure S6**), these areas appeared more integrated within the tissue of hiCHO constructs maintained under static conditions. In contrast, when present in hiCHO constructs exposed to the combined static–dynamic maturation protocol, they were generally confined to the construct periphery and showed limited incorporation into the surrounding cartilage-like matrix.

**Figure 2.**
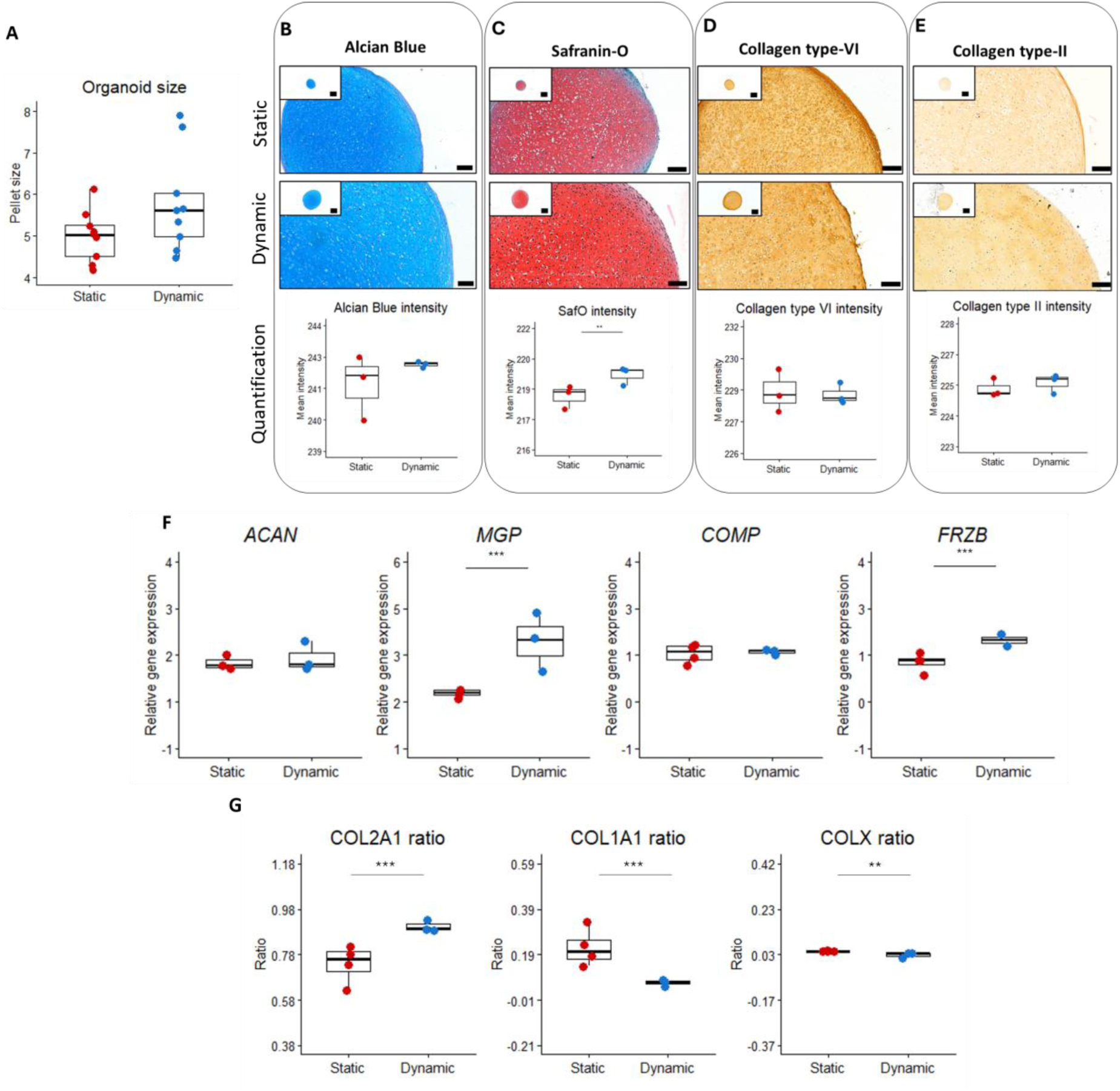
Prolonged maturation of hiCHO constructs under static and dynamic culture conditions. (**A**) Scatter boxplot showing hiCHO organoids size following prolonged maturation in static vs dynamic culture. (**B–E**) Representative histological and immunohistochemical analyses using Alcian Blue, Safranin-O, collagen type VI, and collagen type II staining in W6 hiCHO organoids. Quantification of staining intensity is shown below each representative image. Insets show low and high-magnification overviews (4x, 20x) of the entire constructs. Corresponding images from all biological replicates are shown in **Supplementary Figure S6**. (**F**) Relative mRNA expression levels (−ΔCt) of established chondrogenic markers detected in hiCHOs after 6 weeks maturation in static and dynamic cultures. (**G**) Relative abundance of collagen type II, type I, and type X collagen. Each dot represents an individual construct; bars indicate mean ± SD. Statistical significance was determined using generalized estimating equations (GEE). *P ≤ 0.05, **P ≤ 0.01, ***P ≤ 0.005.

At transcriptional level (**Figure 2F; Supplementary Table S7C),** we showed significant increased *MGP* and *FRZB* expression under the combined static - dynamic culture condition (**Figure 2F**) highlighting ongoing maturation / formation of biomimetic articular cartilage cells. In parallel, the abundance of *COL2A1* relative to *COL1A1* and *COL10A1* was highest in the combined static – bioreactor culture conditions (**Figure 2G; Supplementary Table S7D**). Taken together these findings indicate that hiCHO-made cartilage organoids in both static and dynamic culture systems thrive under a prolonged culture duration, exhibiting continuous deposition of ECM components and progressive acquisition of mature articular cartilage characteristics.

### Integration of dynamically-matured hiCHO organoids in a human OA osteochondral model

Finally, as a proof-of-concept, we explored whether hiCHO organoids generated in dynamic suspension culture conditions, could be retained and integrate within native human cartilage tissue.

To this end, hiCHO-made cartilage organoids matured for 4 weeks of maturation under dynamic suspension cultures were implanted into cartilage defects generated in human preserved cartilage explants (**Figure 3A**), using either a biopsy punch, or a mechanical drill. Prior to implantation, dynamically-matured hiCHO organoids showed robust ECM deposition together with stable cartilage morphology (**Supplementary Figure S7**). Following histological characterization, the hiCHO constructs were manually implanted into the cartilage defects and engrafted explants were harvested after 6 weeks of culturing. Following the culture period, both the untreated drilled and punched defects remained largely empty with little evidence of repair (**Figure 3B-C**). In contrast, as shown by the strong proteoglycan-rich matrix throughout the engrafted regions (**Figure 3D-E**, upper panel), both the treated drilled and the punched defects had retained the hiCHO-made cartilage organoids. Moreover, as shown by higher-magnification Alcian Blue images of the graft-host interface in the defect conditions (**Figure 3D-E**, lower panels, arrowheads) revealed close apposition between the implanted construct and the adjacent cartilage tissue, suggesting progressive integration. Additional collagen type II and collagen type VI staining further confirmed the presence of cartilage-like matrix within the treated defects (**Supplementary Figure S8**). Together, these findings suggest that dynamically matured hiCHO organoids support integration within preserved human cartilage defects *ex vivo*.

**Figure 3.**
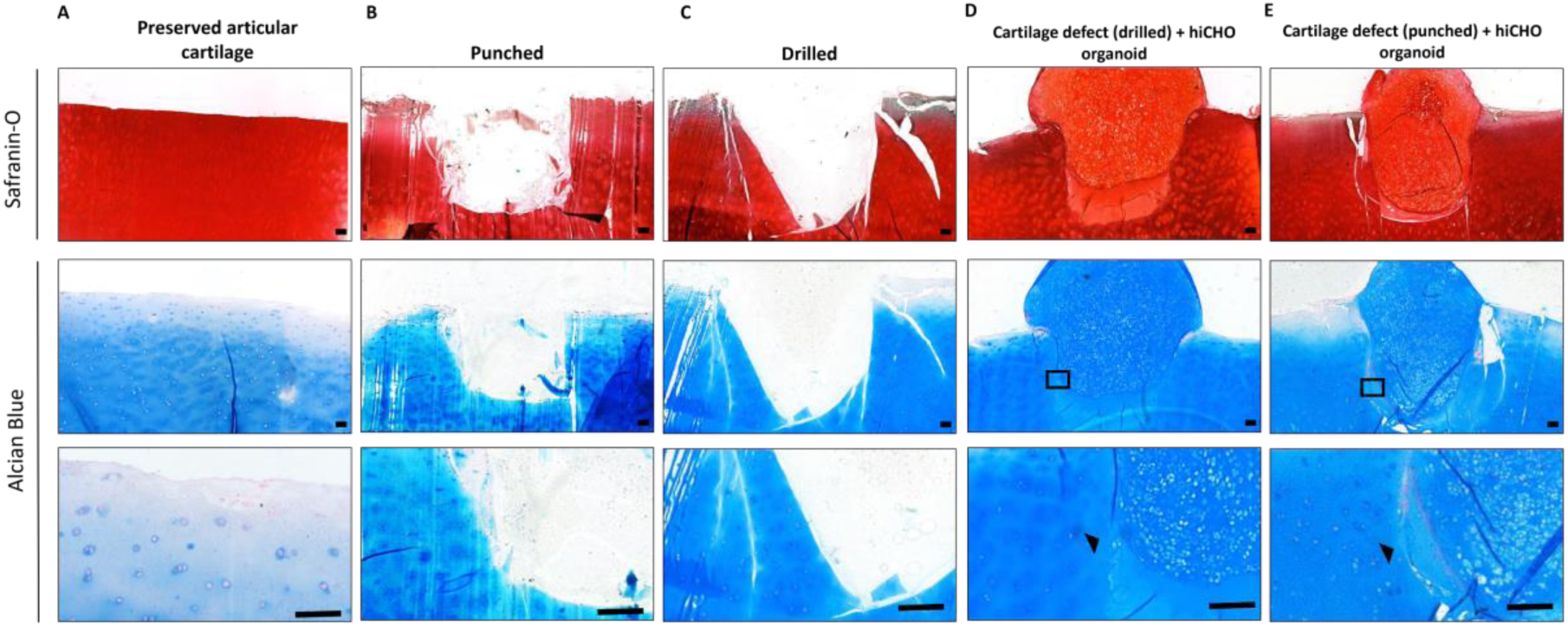
Proof-of-concept assessment of hiCHO-made cartilage organoid integration after 6 weeks in a human cartilage explant model. Representative histological stained sections of *ex vivo* human cartilage explants with and without implanted hiCHO-made cartilage organoids generated under dynamic suspension culture conditions. (**A**) Preserved cartilage control without defect. (**B**) Preserved cartilage with biopsy punched defect without treatment (**C**) Preserved cartilage explant with mechanically drilled defect without treatment. (**D**) Preserved cartilage explant with mechanically drilled defect treated with hiCHO made cartilage organoids. (**E**) . Preserved cartilage explant with biopsy punched defect treated with hiCHO made cartilage organoids. Upper panel Safranin-O staining (4x), middle panel Alcian Blue staining (4x), and lower panel Alcian Blue staining (20x).

## Discussion

In this study we established a workflow that allowed stable hiCPCs maturation into hiCHOs in a dynamic suspension culture condition (CERO bioreactor, OLS). By systematically evaluating inoculation strategy, aggregate preparation, rotation speed, and maturation duration, we identified manually picked pre-formed aggregates cultured at 80 rpm as the most favourable condition for stable hiCHOs maturation. Upon comparing dynamic versus static hiCHO maturation, we demonstrated that dynamic maturation outperformed the standard static condition particularly regarding organoid size, improved tissue organization, and a more favourable chondrogenic cellular phenotype. Importantly, these effects were not accompanied by major differences in proliferative activity, suggesting that dynamic suspension culture primarily supported matrix production and tissue maturation rather than increased cell expansion. In addition, we showed that hiCHO-made cartilage organoids matured under bioreactor conditions homed within human cartilage defects and established matrix continuity with surrounding native tissue, highlighting the translational relevance of this approach. Dynamic suspension culture substantially improved ECM organization. HiCHO organoids matured in the bioreactor showed stronger proteoglycan deposition, more homogeneous matrix distribution, and more clear lacunae formation compared to static cultures.

Interestingly, bioreactor maturation was also associated with a reduced presence of non-cartilage tissue within the hiCHO organoids. Although small regions of such tissue could still be detected after prolonged culture, these were predominantly restricted to the construct periphery and appeared not integrated with the surrounding cartilage matrix than in static cultures. into the cartilage matrix compared hiCHOs matured in static cultures. This may indicate that dynamic suspension culture not only enhances matrix production but also supports more uniform cartilage tissue formation. These histological features resemble aspects of native cartilage organization and are consistent with previous studies showing that bioreactor systems improve nutrient exchange and reduce diffusion gradients within engineered cartilage tissues (10, 11, 13). In parallel, we observed increased *ACAN*, *COMP*, and *MGP* expression together with a more favourable collagen composition profile under dynamic culture conditions. Moreover, we showed that these changes were not accompanied by increased KI67 staining or *KI67* expression, suggesting that the enhanced tissue quality observed under dynamic conditions was primarily driven by matrix production and tissue organization rather than proliferation. Similar findings have been described in mechanically stimulated cartilage cultures, where biomechanical cues predominantly affect ECM synthesis rather than cell expansion . In setting up the dynamic suspension cultures we noted the importance of preserving hiCPC aggregates architecture before transfer to suspension culture. Similar observations have been reported in other stem cell-based chondrogenic systems, where early cell–cell interactions and condensation processes are considered essential for stable cartilage differentiation . We therefore speculate that maintaining early cell–cell interactions together with nascent ECM organization may be crucial during the initial stages of chondrogenesis.

Despite the fact that integration of cartilage implant with surrounding host tissue is considered a major challenge for translational cartilage engineering approaches (23–25), we showed here in a proof-of-concept study that the hiCHO-made cartilage organoids matured in the bioreactor, established ECM continuity with surrounding human cartilage after 6 weeks of *ex vivo* culturing. Although this experiment was limited with N=1 donor and primarily assessed histologically, the observed graft retention and matrix continuity are encouraging findings in the context of cartilage repair. Previous findings using stem cell-derived cartilage organoids have similarly identified graft retention and integration as critical determinants of successful tissue repair (24–26). Our findings therefore support further evaluation of hiCHO-based repair strategies in more advanced and clinically relevant preclinical models.

As evidenced by continuous deposition of ECM components and progressive acquisition of mature articular cartilage characteristics, we showed that hiCHO organoids continued to mature during prolonged culture, even under static conditions. These observations are consistent with previous studies reporting ongoing structural organization and maturation of stem cell-derived cartilage tissues during extended culture period (11, 26, 27). Notably, while prolonged culture under static conditions promoted the formation of prominent chondron-like structures, the introduction of an additional dynamic maturation phase was associated with increased expression of the articular cartilage-associated markers *MGP* and *FRZB* together with a more favourable collagen profile as determined by the normalized ratios of each collagen type over the total collagen expression. These molecular changes are consistent with previous reports describing articular cartilage-like phenotypes in hiPSC-derived cartilage organoids, supporting the notion that dynamic culture promotes progression toward a more mature cartilage state (26–28). Experimental conditions that may further enhance the maturation of hiCHO-derived cartilage formation include optimizing the timing and duration of static and dynamic suspension culture phases, as well as the application of biomechanical stimulation.

Several limitations of the present study should be considered. First, although the bioreactor culture can significantly advance scalable production of off-the-shelf neocartilage constructs for clinical application, the need for manual picking of the hiCPC aggregates remains a limiting time-consuming factor. In addition, although bioreactor culture improved matrix deposition and tissue organization, the mechanisms underlying these effects were not yet determined. Factors such as mechanical loading, nutrient exchange, and oxygen exposure may all contribute to the observed phenotype and warrant further investigation (10, 12, 13). Finally, for the osteochondral explant experiments tissues were obtained from one individual. Additional studies including multiple donor and functional analyses will be necessary to further evaluate long-term tissue integration and stability.

## Conclusions

We established a dynamic suspension culture workflow for maturation of high quality biomimetic hiPSC-derived cartilage organoids in the CERO bioreactor system. Dynamic culture promoted ECM deposition, tissue organization, and cartilage-associated cellular molecular features compared to conventional static culture. Furthermore, hiCHO-derived cartilage organoids generated under dynamic suspension culture conditions successfully integrated into human cartilage defects and established matrix continuity with the surrounding native tissue in a human osteochondral explant model, demonstrating their potential for cartilage repair and regenerative applications.

## List of abbreviations

ACI: Autologous cartilage implantation
ACAN: Aggrecan
BMP-4: Bone morphogenetic protein 4
CDM: Chondrogenic differentiation medium
COL1A1: Collagen type I alpha 1 chain
COL2A1: Collagen type II alpha 1 chain
COL10A1: Collagen type X alpha 1 chain
COMP: Cartilage oligomeric matrix protein
DAB: 3,3′-diaminobenzidine
DMEM: Dulbecco’s Modified Eagle Medium
DMMB: Dimethylmethylene blue
ECM: Extracellular matrix
EDTA: Ethylenediaminetetraacetic acid
FGF-2: Fibroblast growth factor 2
FRZB: Frizzled-related protein
GAPDH: Glyceraldehyde-3-phosphate dehydrogenase
GEE: Generalized estimating equations
GOI: Gene of interest
H&E: Haematoxylin and eosin
hiCHOs: hiPSC-derived chondrocytes
hiCPCs: hiPSC-derived chondroprogenitor cells
hiPSCs: Human induced pluripotent stem cells
IHC: Immunohistochemistry
IMDM: Iscove’s Modified Dulbecco’s Medium
ITS+: Insulin–transferrin–selenium
Ki67: Marker of proliferation Ki67
MGP: Matrix Gla protein
NEAA: Non-essential amino acids
OA: Osteoarthritis
PBS: Phosphate-buffered saline
qPCR: Quantitative polymerase chain reaction
RAAK: Research Arthritis and Articular Cartilage
RNA: Ribonucleic acid
ROI: Region of interest
RT-qPCR: Reverse transcription quantitative polymerase chain reaction
SDHA: Succinate dehydrogenase complex flavoprotein subunit A
sGAG: Sulphated glycosaminoglycan
TGF-β1: Transforming growth factor beta 1
TKR: Total knee replacement

## Declarations

The authors have declared no conflicts of interest.

## Ethics approval and consent to participate

Human osteochondral explants and primary chondrocytes were obtained from osteoarthritis patients undergoing total knee replacement surgery as part of the Research in Articular Osteoarthritis Cartilage (RAAK) study. Ethical approval was granted by the Medical Ethics Committee of the Leiden University Medical Center (P08.239 / P19.013). Written informed consent was obtained from all donors. Approval for generation of hiPSCs from healthy donor skin fibroblasts was obtained under protocol number P13.080.

## Consent for publication

Not applicable.

## Data and materials availability

All data generated or analysed during this study are included in this published article and its supplementary information files. Additional data and materials are available from the corresponding author upon reasonable request.

## Authors’ contribution

Giorgia Mazzini: Collection and/or assembly of data, data analysis and interpretation, manuscript writing, final approval of manuscript.

Evelyn Houtman: Collection and/or assembly of data, final approval of manuscript

Marcella van Hoolwerff: Collection and/or assembly of data, final approval of manuscript

Merel Janssen: Collection and/or assembly of data, final approval of manuscript.

Sana Sayedipour: Collection and/or assembly of data, final approval of manuscript.

Ghazaleh Hajmousa: Collection and/or assembly of data, final approval of manuscript.

Roxanne E. Kieltyka: Collection and/or assembly of data, final approval of manuscript.

Rachid R. Mahdad: Collection and/or assembly of data, final approval of manuscript.

Yolande F.M. Ramos: Conception and design, data analysis and interpretation, manuscript writing, final approval of manuscript.

Ingrid Meulenbelt: Conception and design, data analysis and interpretation, manuscript writing, final approval of manuscript.

## Acknowledgements

We thank all study participants of the RAAK study. The Leiden University Medical Centre and Alrijne Leiderdorp have and are supporting the RAAK. For that matter we thank Rachid Mahdad (Alrijne Leiderdorp) Enrike van der Linden, and Anika Rabelink-Hoogenstraaten (LUMC) for their contribution to the collection of the joint tissues. Data is generated within the scope of the Medical Delta programs Regenerative Medicine 4D: Generating complex tissues with stem cells and printing technology and Improving Mobility with Technology.

