## Supplemental materials for "Dynamic suspension culture enhances scalable maturation of hiPSC-derived cartilage organoids for regenerative medicine"

This document contains:

**Supplementary Figure S1-S8**

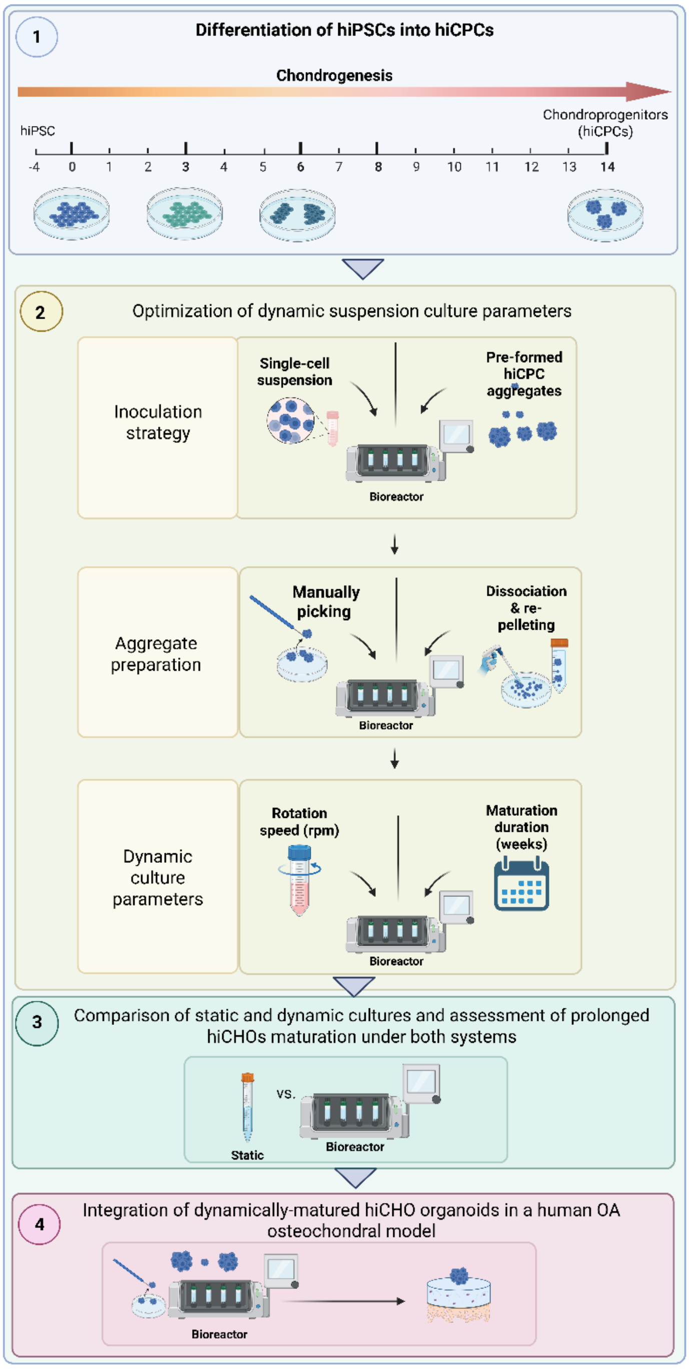


**Supplementary Figure S1. Schematic overview of the experimental workflow.** Human induced pluripotent stem cells (hiPSCs) were differentiated into chondroprogenitor cells (hiCPCs) over 14 days. Subsequently, dynamic suspension culture conditions were optimized by evaluating inoculation strategy, aggregate preparation method, rotation speed, and maturation duration. Optimized dynamic suspension culture conditions were then compared with conventional static culture for hiCHO maturation. Finally, dynamically matured hiCHO constructs were evaluated for integration in a human osteochondral explant model.

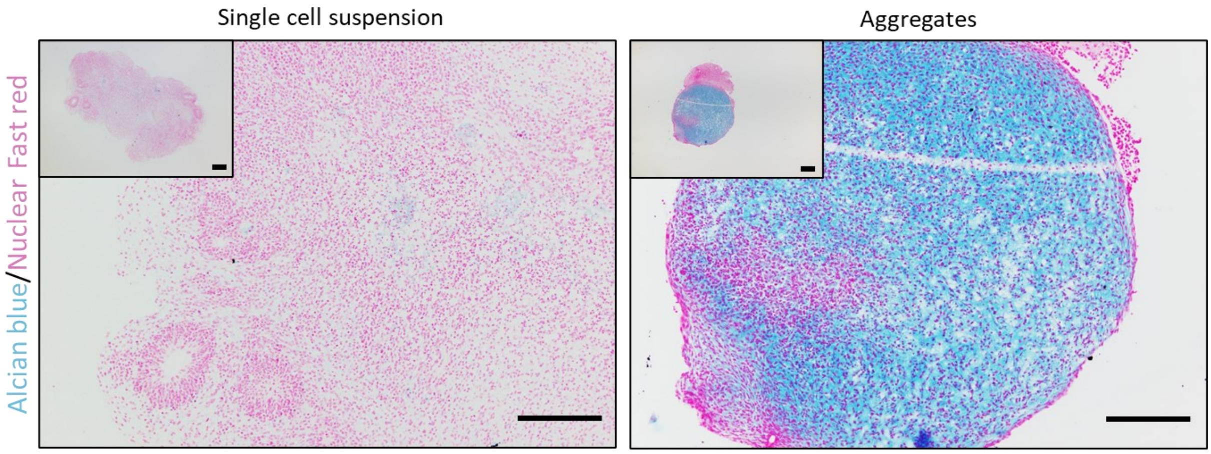


**Supplementary Figure S2. Comparison of aggregate formation strategies for dynamic suspension culture.** Representative Alcian Blue/Nuclear Fast Red staining of organoids generated from day-14 hiCPC aggregates introduced into dynamic suspension culture either as single-cell suspensions or as pre-formed aggregates. Insets show low and high-magnification overviews (4x, 10x) of the corresponding constructs.

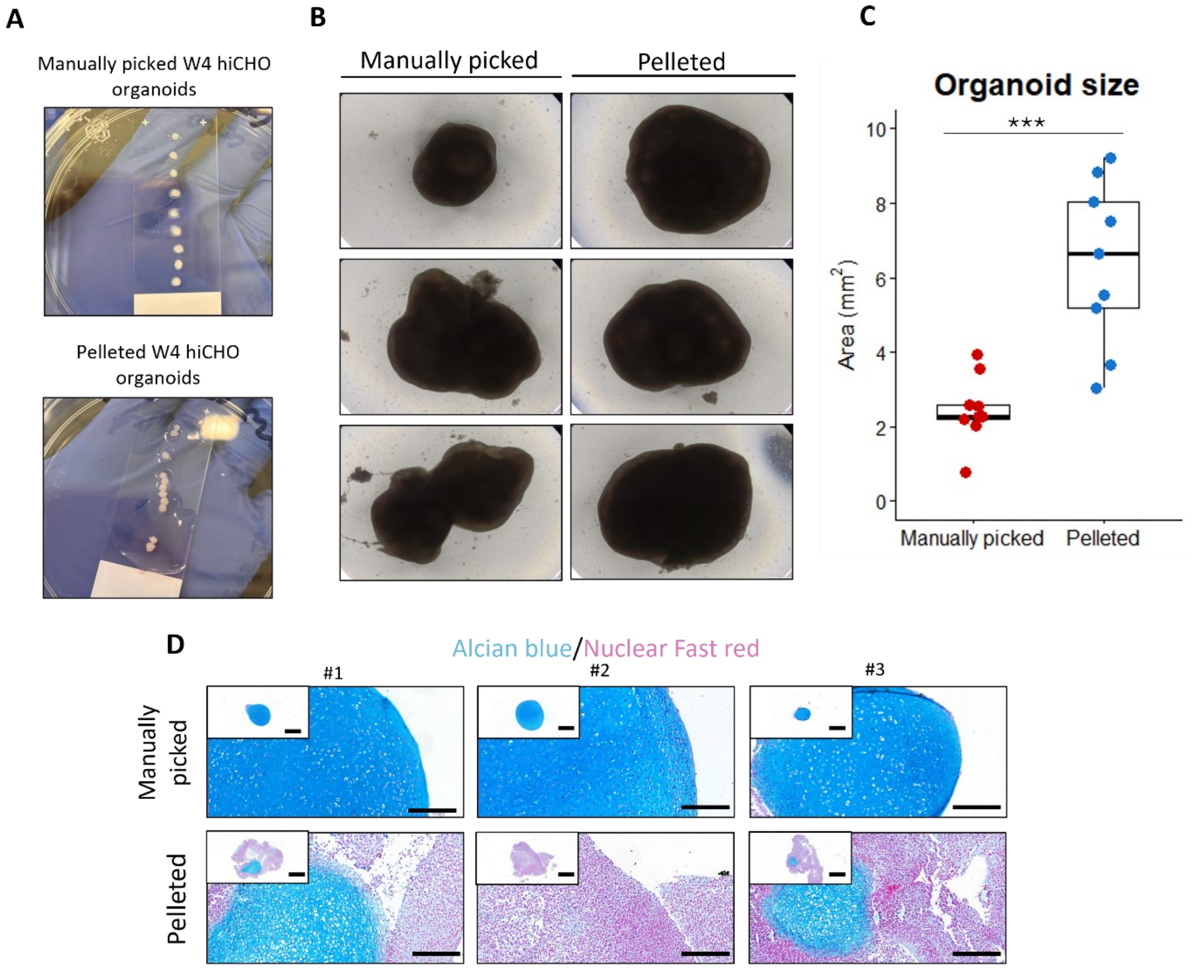


**Supplementary Figure S3. Comparison of W4 manually picked and pelleted hiCHO organoids matured in the bioreactor.** (**A**) Representative macroscopically images of manually picked and pelleted hiCHO organoids after 4 weeks of maturation. (**B**) Representative brightfield images of individual constructs generated from manually picked or pelleted hiCHO organoids after 4 weeks of maturation in the bioreactor. (**C**) Quantification of construct cross-sectional area. (**D**) Representative Alcian Blue/Nuclear Fast Red staining of three independent constructs generated from manually picked or pelleted hiCHO organoids after 4 weeks in the bioreactor. Insets show low and high-magnification overviews (4x, 20x) of the corresponding constructs. Statistical significance was determined using GEE analysis. ***P ≤ 0.005.

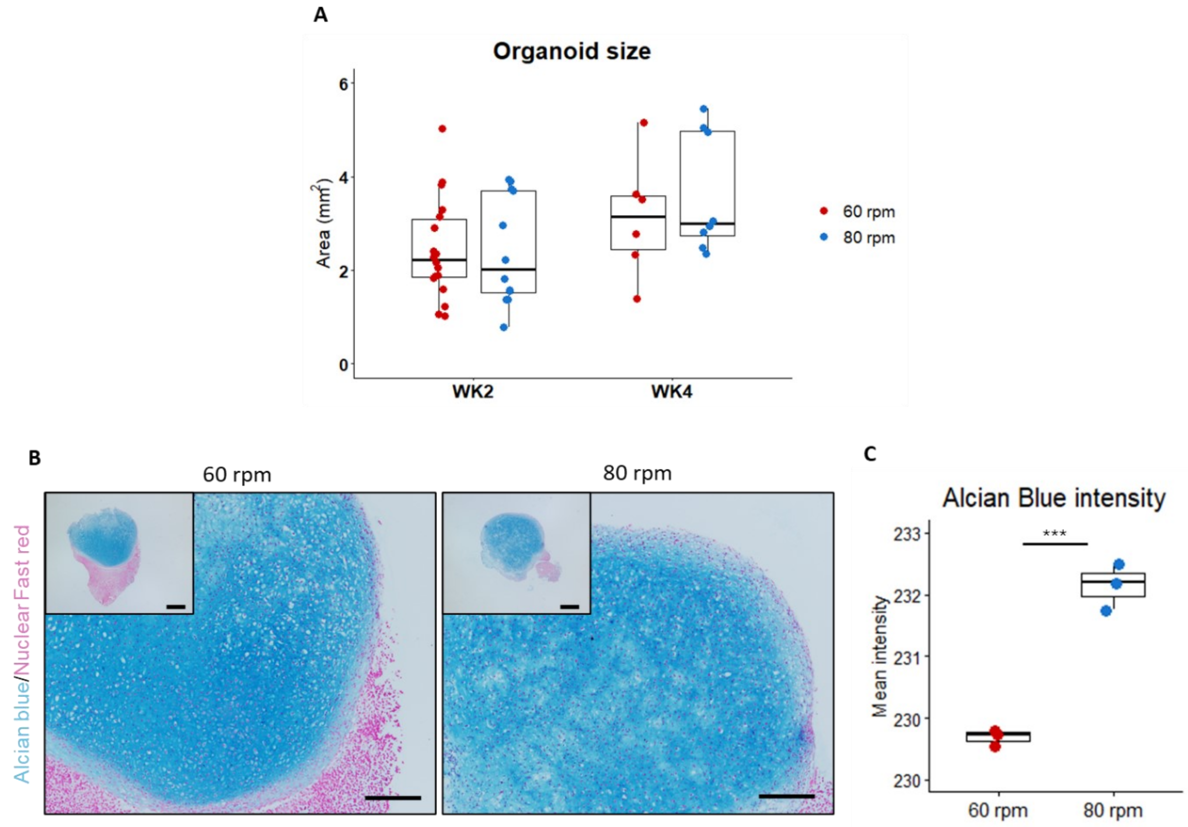


**Supplementary Figure S4. Optimization of dynamic suspension culture conditions for hiCHO maturation.** (**A**) Pellet size measurement of hiCHO constructs cultured under dynamic suspension conditions (bioreactor) at 60 or 80 rpm for 2 or 4 weeks. (**B**) Representative Alcian Blue and collagen type II staining of constructs matured for 4 weeks at 60 or 80 rpm. Insets show low and high-magnification overviews (4x, 10x) of the corresponding constructs. (**C**) Quantification of Alcian Blue staining intensity. Individual measurements are shown with mean ± SD. Statistical significance was determined using GEE analysis. ***P ≤ 0.005.

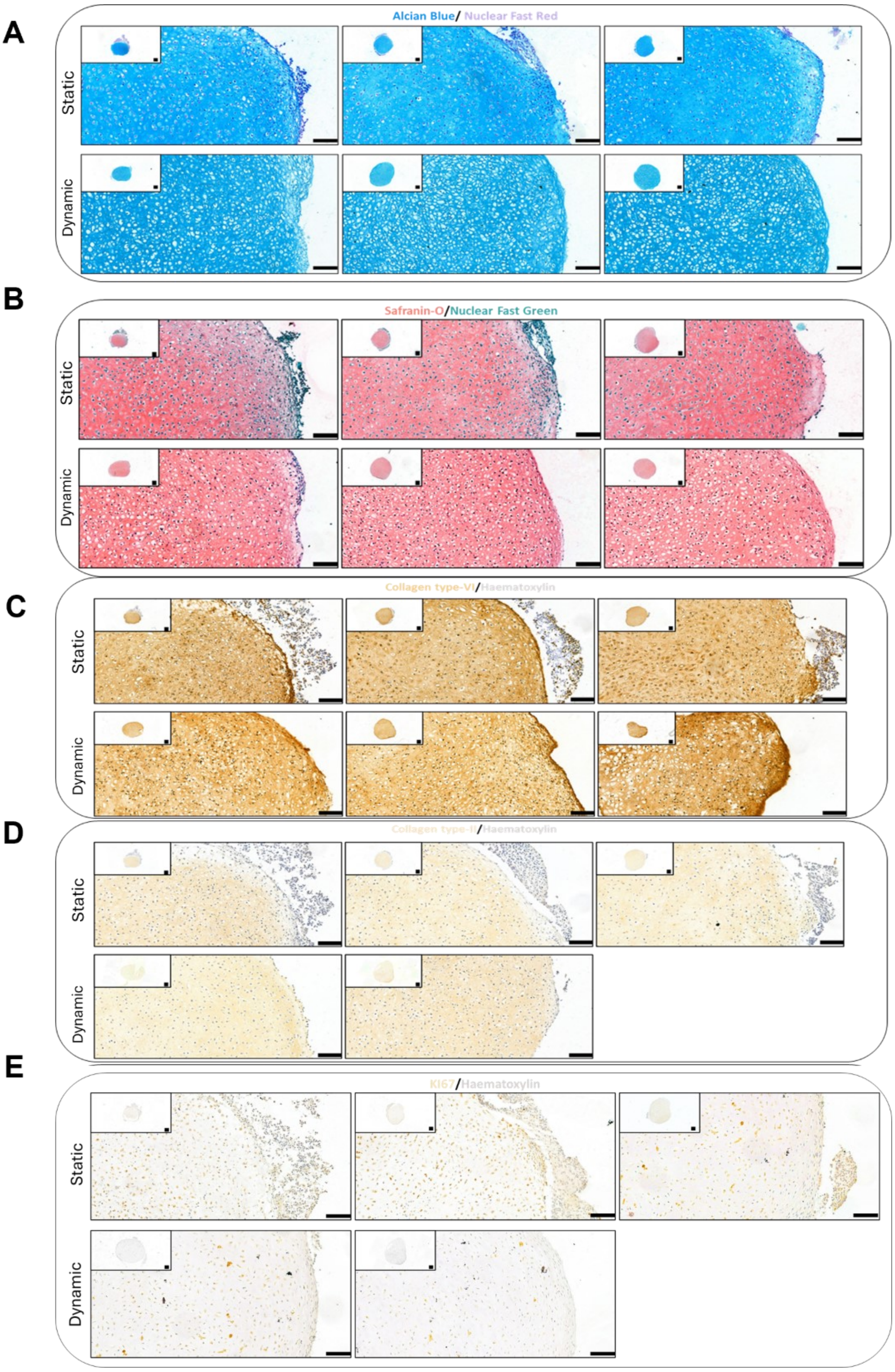


**Supplementary Figure S5. Histological and immunohistochemical analyses of all biological replicates included in the 4-week maturation experiment**. Representative sections of hiCHO constructs matured for 4 weeks under static or dynamic suspension culture conditions. (**A**) Alcian Blue/Nuclear Fast Red staining. (**B**) Safranin-O/Fast Green staining. (**C**) Collagen type VI immunostaining. (**D**) Collagen type II immunostaining. (**E**) Ki67 immunostaining. Insets show low and high-magnification overviews (4x,20x) of the corresponding hiCHO constructs. Quantification of staining intensity is presented in **Figure 1** and **Supplementary Table S8**.

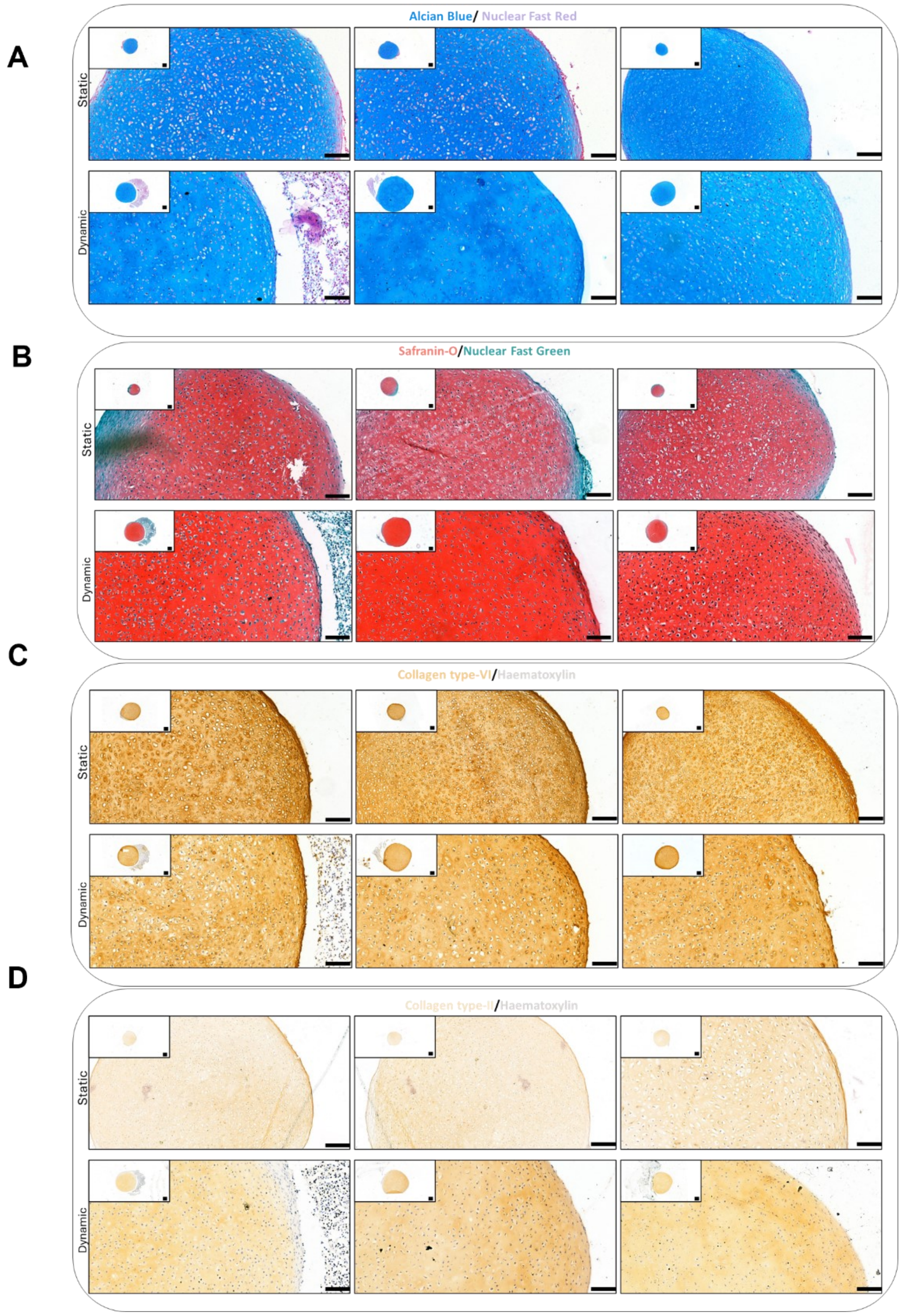


**Supplementary Figure S6. Histological and immunohistochemical analyses of all biological replicates included in the prolonged maturation experiment.** Representative sections of hiCHO constructs matured for 6 weeks under either continuous static culture conditions or sequential static and dynamic suspension culture conditions. (**A**) Alcian Blue/Nuclear Fast Red staining. (**B**) Safranin-O/Fast Green staining. (**C**) Collagen type VI immunostaining. (**D**) Collagen type II immunostaining. Insets show low and high-magnification overviews (4x, 20x) of the corresponding hiCHO constructs. Quantification of staining intensity is presented in Figure 2 and Supplementary Table S10.

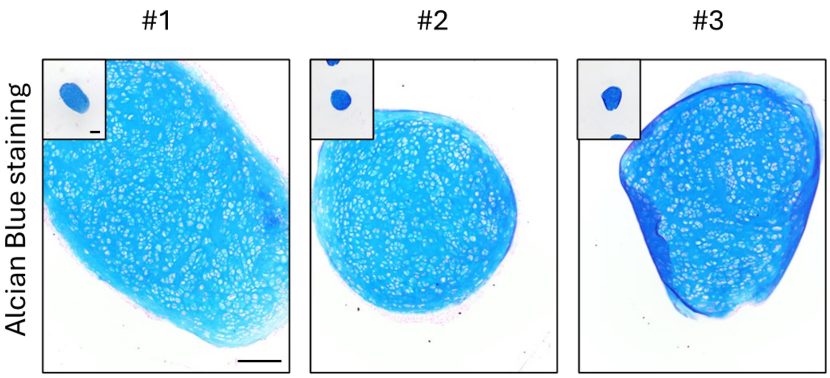


**Supplementary Figure S7. Histological characterization of hiCHO constructs before (W4) implantation.** Representative Alcian Blue/Nuclear Fast Red staining of dynamically matured hiCHO constructs prior to implantation in the osteochondral explant model. Insets show low and high-magnification overviews (4x, 10x) of the corresponding constructs.

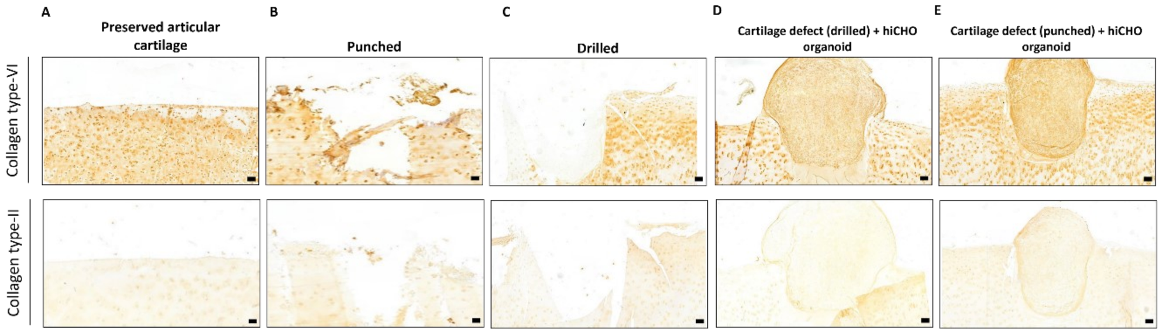


**Supplementary Figure S8. Additional characterization of cartilage matrix markers in the human osteochondral explant model.** Representative immunohistochemical staining of ex vivo human cartilage explants with and without implanted hiCHO constructs generated under dynamic suspension culture conditions. (**A**) Preserved articular cartilage control without defect. (**B**) Preserved cartilage with a biopsy-punched defect without treatment. (**C**) Preserved cartilage explant with a mechanically drilled defect without treatment. (**D**) Preserved cartilage explant with a mechanically drilled defect treated with a hiCHO construct. (**E**) Preserved cartilage explant with a biopsy-punched defect treated with a hiCHO construct. Upper panel: collagen type VI immunostaining. Lower panel: collagen type II immunostaining. Images were acquired at 4× magnification.
