## Supplemental tables for "Dynamic suspension culture enhances scalable maturation of hiPSC-derived cartilage organoids for regenerative medicine"

This document contains:

**Supplementary Tables S1-S8**

**Supplementary Table S1.** Parameters for dynamic suspension culture using the CERO 3D bioreactor.

| Parameter | Inoculation phase | Culture phase |
|---|---|---|
| Rotation period | 1 s | 1 s |
| Rotation speed | 80 rpm | 80 rpm |
| Rotation pause | 2 s | 2 s |
| Condition | Normoxia | Hypoxia |
| Protocol duration | 72 h | ∞ |

**Supplementary Table S2.** Primer sequences to measure mRNA expression levels

| Region of interest | Forward primer (5' --> 3') | Reverse primer (5' --> 3') |
|---|---|---|
| *ACAN* | AGAGACTCACACAGTCGAAACAGC | CTATGTTACAGTGCTCGCCAGTG |
| *COL1A1* | GTGCTAAAGGTGCCAATGGT | ACCAGGTTCACCGCTGTTAC |
| *COL2A1* | CTACCCCAATCCAGCAAACGT | AGGTGATGTTCTGGGAGCCTT |
| *COL10A1* | GGCAACAGCATTATGACCCA | TGAGATCGATGATGGCACTCC |
| *COMP* | ACAATGACGGAGTCCCTGAC | TCTGCATCAAAGTCGTCCTG |
| *FRZB* | ATTGACTTCCAGCACGAGCC | CGAGTGGCGGTACTTGATGAG |
| *GAPDH* | TGCCATGTAGACCCCTTGAAG | ATGGTACATGACAAGGTGCGG |
| *KI67* | ACACATTGTCCTCAGCCTTCT | GTGCCTGAGATCAAGAAAGACA |
| *MGP* | CGCCCCCAGATTGATAAGTA | TCTCCTTTGACCCTCACTGC |
| *SDHA* | TGGGAACAAGAGGGCATCTG | GCCTACCACCACTGCATCAA |

**Supplementary Table S3.** The Primary and Secondary antibodies used for immunohistochemical (IHC) analysis and their concentrations in PBS

| Detection of | Primary antibody | Secondary antibody |
|---|---|---|
| Collagen type-II | Rabbit anti-collagen type-II (abcam34712; Abcam), 1:250 | Goat anti-rabbit 488 (ab150077, Abcam), 1:500 |
| Collagen type-VI | Rabbit anti-collagen type-VI (abcam6588; Abcam), 1:250 | Goat anti-rabbit 488 (ab150077, Abcam), 1:500 |
| KI67 | Rabbit anti-KI67 (abcam92742; Abcam), 1:500 | Goat anti-rabbit 488 (ab150077, Abcam), 1:500 |

**Supplementary Table S4**. Statistical difference in pellet size between W4 manually picked and pelleted hiCHO organoids, determined by GEE (reference group: Manually picked).

| Comparison | B | Std. Error | 95% Wald Confidence Interval | | Wald Chi-Square | Sig. |
|---|---|---|---|---|---|---|
| Manually picked vs Pelleted | 3,95 | 0,62 | 2,74 | 5,16 | 41,17 | **1.39e-10** |

**Supplementary Table S5A.** Statistical differences in organoid size (mm²) of hiCHO-derived neo-cartilage cultured under dynamic conditions (60 and 80 rpm and week 2 & 4), determined by generalized estimating equations (GEE).

| Effect | Comparison | B | Std. Error | 95% Wald Confidence Interval | | Wald Chi-Square | Sig. |
|---|---|---|---|---|---|---|---|
| Time effect | Week 4 vs Week 2 within 60 rpm | 0,7 | 0,54 | -0,36 | 1,75 | 1,68 | 1.95e-01 |
| Time effect | Week 4 vs Week 2 within 80 rpm | 1,23 | 0,53 | 0,19 | 2,27 | 5,33 | **2.09e-02** |
| RPM effect | 80 rpm vs 60 rpm at Week 2 | -0,02 | 0,4 | -0,82 | 0,77 | 0 | 9.52e-01 |
| RPM effect | 80 rpm vs 60 rpm at Week 4 | 0,51 | 0,64 | -0,75 | 1,76 | 0,63 | 4.29e-01 |
| Interaction | Time × RPM | 0,53 | 0,76 | -0,95 | 2,01 | 0,49 | 4.83e-01 |

Data were analyzed using generalized estimating equations (GEE). B, regression coefficient; SE, standard error; [illegible] interval; Wald $\chi^2$, Wald chi-square statistic.

**Supplementary Table S5B.** Statistical differences in hiCHO organoids staining intensity between 60 rpm and 80 rpm dynamic culture conditions at Week 4, determined by GEE.

| Comparison | B | Std. Error | 95% Wald Confidence Interval | | Wald Chi-Square | Sig. |
|---|---|---|---|---|---|---|
| Alcian Blue | 80 rpm vs 60 rpm | 1,84 | 0,14 | 1,46 | 2,23 | **1.84e-04** |

**Supplementary Table S6A.** Statistical difference in organoid size (mm²) between manually picked static vs dynamic W4 hiCHO derived neo-cartilage organoids, determined by GEE. Reference group: Static

| Comparison | B | Std. Error | 95% Wald Confidence Interval | | Wald Chi-Square | Sig. |
|---|---|---|---|---|---|---|
| Static vs CERO | 2,37 | 0,39 | 1,6 | 3,13 | 36,83 | **1.29e-09** |

**Supplementary Table S6B.** Statistical difference in sGAGs/DNA ratio between W4 manually picked static vs dynamic W4 hiCHO derived neo-cartilage organoids, determined by GEE. Reference group: Static

| Comparison | B | Std. Error | 95% Wald Confidence Interval | | Wald Chi-Square | Sig. |
|---|---|---|---|---|---|---|
| Manually picked static vs CERO | 0,64 | 0,23 | 0,19 | 1,09 | 7,8 | **5.24e-03** |

**Supplementary Table S6C.** Statistical difference in hiCHO organoids staining intensity between manually picked static vs dynamic hiCHO derived neo-cartilage organoids, determined by GEE. Reference group: Static

| Comparison | Condition | B | Std. Error | 95% Wald Confidence Interval | | Wald Chi-Square | Sig. |
|---|---|---|---|---|---|---|---|
| Static vs CERO | Alcian Blue | 0,75 | 0,32 | 0,12 | 1,39 | 5,39 | 2.03e-02 |
| | Collagen type II | -0,12 | 0,17 | -0,2 | 0,45 | 0,56 | 4.55e-01 |
| | Collagen type VI | 3,96 | 0,69 | -5,32 | -2,6 | 32,56 | **1.15e-08** |
| | Ki67 | -0,1 | 0,22 | -0,53 | 0,33 | 0,19 | 6.59e-01 |
| | Safranin-O | 4,77 | 0,94 | -6,62 | -2,93 | 25,78 | **3.82e-07** |

**Supplementary Table S6D**. Statistical difference in gene expression (minDCT) of manually picked static and dynamic W4 hiCHO derived neo-cartilage organoids, determined by GEE (reference group: Static)

| Comparison | Gene | B | Std. Error | 95% Wald Confidence Interval | | Wald Chi-Square | Sig. |
|---|---|---|---|---|---|---|---|
| Static vs CERO | *COL2A1* | 4,36 | 0,2 | 3,97 | 4,75 | 474,27 | **3.8e-105** |
| | *ACAN* | 3,21 | 0,15 | 2,91 | 3,52 | 432,5 | **4.6e-96** |
| | *COMP* | 4,15 | 0,73 | 2,73 | 5,57 | 32,7 | **1.1e-08** |
| | *COL10A1* | 2,21 | 1,12 | 0,02 | 4,41 | 3,9 | 4.8e-02 |
| | *COL1A1* | -1,27 | 0,56 | -2,38 | -0,17 | 5,12 | 2.4e-02 |
| | *MGP* | 4,41 | 0,31 | 3,81 | 5,01 | 205,6 | **1.3e-46** |
| | *FRZB* | -1,9 | 0,27 | -2,43 | -1,37 | 48,51 | **3.3e-12** |
| | *KI67* | 1,14 | 0,63 | -0,09 | 2,37 | 3,32 | 6.8e-02 |

**Supplementary Table S6E**. Statistical differences in matrix composition ratios between manually picked static and dynamic W4 hiCHO derived neo-cartilage organoids, determined by GEE (reference group: Static)

| Comparison | Gene | B | Std. Error | 95% Wald Confidence Interval | | Wald Chi-Square | Sig. |
|---|---|---|---|---|---|---|---|
| Static vs CERO | *COL2A1/(COL1A1+COL2A1+COL10A1)* ratio | 0,23 | 0,05 | 0,13 | 0,33 | 20,35 | **6.5e-06** |
| | *COL1A1/(COL2A1+COL2A1+COL10A1)* ratio | -0,19 | 0,02 | -0,23 | -0,15 | 81,96 | **1.4e-19** |
| | *COL10A1/(COL2A1+COL1A1+COL10A1)* ratio | -0,03 | 0,04 | -0,12 | 0,05 | 0,64 | 4.2e-01 |

**Supplementary Table S7A.** Statistical difference in size between manually picked static vs dynamic W6 hiCHO neo-cartilage organoids, determined by GEE. Reference group: Static

| Comparison | B | Std. Error | 95% Wald Confidence Interval | | Wald Chi Square | Sig. |
|---|---|---|---|---|---|---|
| Static vs CERO | 0,81 | 0,43 | -0,03 | 1,64 | 3,59 | 5.80e-02 |

**Supplementary Table S7B.** Statistical difference in staining intensity between manually picked static vs dynamic W6 hiCHO derived neo-cartilage organoids, determined by GEE. Reference group: Static

| Comparison | Condition | B | Std. Error | 95% Wald Confidence Interval | | Wald Chi-Square | Sig. |
|---|---|---|---|---|---|---|---|
| Static vs CERO | Alcian Blue | 0,87 | 0,66 | -0,44 | 2,17 | 1,7 | 1.92e-01 |
| | Collagen type II | 0,25 | 0,28 | -0,29 | 0,79 | 0,81 | 3.67e-01 |
| | Collagen type VI | 0,14 | 0,49 | -0,82 | 1,09 | 0,08 | 7.80e-01 |
| | Safranin-O | 0,94 | 0,32 | 0,32 | 1,56 | 8,9 | **2.85e-03** |

**Supplementary Table S7C.** Statistical difference in gene expression (minDCT) of manually picked static vs dynamic W6 hiCHO derived neo-cartilage organoids, determined by GEE. Reference group: Static

| Comparison | Gene | B | Std. Error | 95% Wald Confidence Interval | | Wald Chi-Square | Sig. |
|---|---|---|---|---|---|---|---|
| Static vs CERO | *ACAN* | 0,13 | 0,21 | -0,28 | 0,54 | 0,37 | 5.45e-01 |
| | *COL10A1* | 0,68 | 0,13 | 0,41 | 0,94 | 25,33 | **4.83e-07** |
| | *COL1A1* | -0,06 | 0,21 | -0,47 | 0,35 | 0,08 | 7.72e-01 |
| | *COL2A1* | 1,88 | 0,31 | 1,26 | 2,49 | 35,63 | **2.39e-09** |
| | *COMP* | 0,05 | 0,12 | -0,19 | 0,29 | 0,17 | 6.81e-01 |
| | *FRZB* | 1,38 | 0,37 | 0,65 | 2,11 | 13,62 | **2.24e-04** |
| | *MGP* | 0,59 | 0,15 | 0,3 | 0,88 | 16,15 | **5.86e-05** |

**Supplementary Table S7D.** Statistical difference in collagen ratios between manually picked static vs dynamic W6 hiCHO derived neo-cartilage organoids, determined by GEE. Reference group: Static

| Comparison | Gene | B | Std. Error | 95% Wald Confidence Interval | | Wald Chi-Square | Sig. |
|---|---|---|---|---|---|---|---|
| Static vs CERO | *COL2A1/(COL1A1+COL2A1+COL10A1)* ratio | 0,16 | 0,04 | 0,09 | 0,24 | 18,62 | **1.59e-05** |
| | *COL1A1/(COL2A1+COL2A1+COL10A1)* ratio | -0,15 | 0,04 | -0,22 | -0,08 | 17,89 | **2.34e-05** |
| | *COL10A1/(COL2A1+COL1A1+COL10A1)* ratio | -0,02 | 0,01 | -0,03 | -0,01 | 9,38 | **2.20e-03** |

**Supplementary Table S8.** Patient characteristics of knee joint used to isolate osteochondral explants for the study **(Figure 3)**

| Parameters | Donor |
|---|---|
| Sex | Female |
| Age | 80 |
| KL-score | 4 |
